# Non-responsive Celiac disease symptoms associated with microbiome network structure and function

**DOI:** 10.1101/2023.10.14.562350

**Authors:** Laura Judith Marcos-Zambrano, Blanca Lacruz-Plegezuelos, Alberto Valdés, Elena Aguilar-Aguilar, Viviana Loria-Kohen, Alejandro Cifuentes, Ana Ramírez de Molina, Enrique Carrillo de Santa Pau

**Author notes:** Address correspondence to Laura Judith Marcos-Zambrano. Viviana Loria-Kohen, Departamento de Nutrición y Ciencia de los Alimentos, Faculty of Pharmacy, Universidad Complutense de Madrid, Madrid, Spain.

## Abstract

Non-responsive celiac disease (NRCD) poses a challenge for clinicians due to the persistence of symptoms despite maintaining a gluten-free diet (GFD). This study investigated the gut microbiome, mucosal integrity, and metabolomic profiles of 39 NRCD patients to gain insights into the underlying mechanisms contributing to symptom persistence. Two distinct clusters of patients were identified based on clinical and demographic variables not influenced by gluten consumption.

Cluster 1, labelled “Low-grade symptoms,” displayed milder symptoms and lower inflammatory markers. In contrast, Cluster 2, named “High-grade symptoms,” exhibited more severe gastrointestinal and extraintestinal symptoms, along with elevated inflammatory markers and increased intestinal permeability.

Despite similar mucosal damage in both clusters, network analysis of the gut microbiome revealed specific microbial taxa with potential functional implications. Cluster 1 displayed a microbiome associated with immune homeostasis and gut barrier integrity, potentially lowering inflammation and symptom severity. In contrast, Cluster 2 had a distinct microbiome linked to lactate production, Th17 activation, possibly contributing to heightened inflammation and gastrointestinal symptoms.

Metabolomic analysis revealed differential metabolites between clusters, particularly in amino acid metabolism pathways. Metabolites associated with specific symptoms were identified, implicating their potential role in symptom manifestation. Notably, vitamin D deficiency was observed in both clusters, suggesting its relevance in the context of NRCD.

The study highlights the importance of the gut microbiome, mucosal integrity, and metabolic pathways in symptom persistence among NRCD patients. The associations between microbial-derived metabolites and symptom severity provide valuable insights into potential therapeutic targets. Further research is needed to validate these findings and develop targeted interventions for improving clinical outcomes in NRCD patients.

## Background

Celiac disease (CD) is a multifaceted autoimmune disorder that arises as a response to the consumption of gluten, the primary protein stored in wheat, barley, and rye (Caio et al., 2019). This condition primarily affects individuals with a genetic predisposition and leads to increased levels of autoantibodies specific to celiac disease. Consequently, it manifests as varying degrees of inflammation in the small intestine and gives rise to a broad spectrum of gastrointestinal and extra-intestinal symptoms (Catassi et al., 2022).

Currently, the only effective approach for managing CD involves strict and lifelong compliance with a gluten-free diet (GFD) (Al-Toma et al., 2019). Typically, this course of action leads to the alleviation of inflammation in the small intestine (Caio et al., 2019). Despite a significant improvement observed in most individuals with CD shortly after eliminating gluten from their diet, a notable percentage ranging from 7% to 30% continue to experience symptoms or exhibit clinical indications suggestive of CD, despite adhering to a GFD (Leonard et al., 2017). This clinical challenge, encompassing a diverse array of diagnoses, is referred to as nonresponsive celiac disease (NRCD) (Sansotta et al., 2018).

NRCD can be further categorized as primary if there is an initial lack of response to a GFD, or secondary if signs, symptoms, or laboratory abnormalities consistent with CD reoccur following an initial period of normalization while maintaining GFD (Leffler et al., 2007). In previous studies, the primary contributor to NRCD was unintentional ingestion of gluten, accounting for approximately 50% of cases (Abdulkarim et al., 2002; Leffler et al., 2007). However, the aetiologies of NRCD exhibit significant variability and can include lymphoma, small-intestinal bacterial overgrowth (SIBO), microscopic colitis, pancreatic insufficiency, disaccharidase deficiency, and irritable bowel syndrome (IBS) (Leffler et al., 2007; O’Mahony et al., 1996; Penny et al., 2020).

Current investigations are focused on the intestinal microbiota, which is known to influence the immune regulation in the gut and is involved in health and disease, not only affecting the gut (Levy et al., 2017). It is well known that patients with CD show a dysbiosis, with a decrease in species from genus *Bifidobacterium* and an overrepresentation of proinflammatory groups, a picture that is likely to precede the onset of the disease and persist after diagnosis, at least in some cases (Arcila-Galvis et al., 2022). Considering that diet is one of the major drivers of human gut microbiota composition (Dahl et al., 2020), it is reasonable to think that GFD can directly affect the host’s physiology and metabolism, as well as gut bacterial activity. For this reason, the interactions between the host’s genetics, GFD, and microbiota need to be deepened. Noteworthy, nutritional regimens, including GFD, involve the production of specific microbial and human metabolites (Bascuñán et al., 2020).

As NRCD encompasses several potentially severe disorders with diverse treatment options and prognoses, ensuring efficient and cost-effective management for patients affected by this syndrome can present challenges. In the present work, we aim to evaluate the persistence of symptoms in NRCD with the gut microbiome and metabolome.

## Methods

### Participants recruitment

A purposive sampling strategy was used to engage participants. Inclusion criteria were Men and women aged between 18 and 65 years, agree to participate in the study voluntarily and give written informed consent, have CD confirmed by intestinal biopsy or documented medical diagnosis at least 12 months before the study, and be following a GFD for at least 12 months. Presence of any of the following symptoms: diarrhoea, soft stools or constipation, abdominal pain, bloating, nausea, discomfort from noise and bowel movements, and tenesmus. Exclusion criteria were Individuals with other pathologies of the digestive system (Crohn’s disease, irritable bowel syndrome, ulcerative colitis, colon cancer), individuals who have undergone digestive system surgery (e.g., short bowel syndrome). Subjects with serious diseases (liver, kidney, cancer) or autoimmune diseases or who use immunosuppressive or anti-inflammatory drugs. Subjects with dementia, mental illness or decreased cognitive function. Subjects usually treated with probiotics or taking antibiotics. Pregnant or breastfeeding women, or any other condition that limits compliance with procedures established in the protocol, as well as adherence to treatment by the subject.

### Ethical Aspects and Data Processing

Protocols and methodology used in the present study comply with the ethical principles for research involving human subjects laid down in the Declaration of Helsinki (1964) and its modifications. The study was approved by the Research Ethics Committee of the IMDEA Food Foundation (PI-032 Approval date: June 12th, 2017). Participants were informed in detail about the different stages of the project both orally and in writing. The researchers collected signed informed consent prior to the first evaluation. This document included a specific consent to microbial profiling. Data compiled along the study was processed by applying dissociation criteria, making the volunteers’ data anonymous, in compliance with the current Spanish legislation (Organic Law 15/1999 of December 13th, on the protection of Personal Data).

### Symptomatology and quality of life assessment

Symptomatology data was recorded through validated questionnaires: (i) the Gastrointestinal Symptom Rating Scale Celiac Disease (GSRS) (Svedlund et al., 1988), a validated 15-item questionnaire employed to assess the severity of gastrointestinal symptoms associated with CD and other related gastrointestinal conditions. The survey comprises five distinct sub-dimensions: Indigestion, diarrhoea, abdominal pain, reflux, and constipation. Scoring for each sub-dimension involves calculating the mean value of the respective items, while the overall GSRS-CD score is determined by computing the mean value across all 15 items. The scoring is performed on an 11-point Likert scale, ranging from 0 to 10, where higher scores indicate more pronounced symptom severity. (ii) Celiac Disease Patient-Reported Outcome (Leffler et al., 2015) is a validated 9-item questionnaire used to evaluate the severity of gastrointestinal and extraintestinal symptoms from CD. Responses are scored on an 11-point (0–10) Likert scale, with higher scores indicating greater symptom severity.

### Dietary intake and gluten-free diet compliance

A validated food record of 72h consumption (Ortega et al., 2015) in which participants had to sum up all food and drinks ingested during three days (2 weekdays and 1 Sunday or holiday) was delivered. Afterwards, the data were tabulated and analyzed by a certified nutritionist using the DIAL nutritional software v 3.15 (Alce Ingeniería, Madrid, Spain) in order to obtain information about macro and micronutrients. Gluten-free diet compliance was monitored by the celiac dietary adherence test (CDAT) (Fueyo-Díaz et al., 2015), a validated questionnaire that considers five crucial dimensions pertaining to compliance with a GFD: the presence of CD symptoms, the patient’s understanding of the disease and its treatment, confidence in the effectiveness of the treatment, motivating factors for adhering to a GFD, and self-reported adherence to the diet. The questionnaire comprises 7 items, each rated on a 5-point Likert scale, resulting in a total score ranging from 7 to 35 points. The scoring interpretation is as follows: a score of 7 points indicates excellent GFD adherence; 8–12 points suggest very good GFD adherence; 13–17 points imply insufficient or inadequate GFD adherence and scores exceeding 17 points indicate poor GFD adherence.

### Anthropometric and Metabolic Measurements

Anthropometric measurements were determined early in the morning, by previously trained nutritionists, following standardized protocols. Height was determined using a Leicester height rod with millimetric accuracy (Biological Medical Technology SL, Barcelona, Spain). Body weight, fat mass (FM) percentage, and muscle mass (MM) percentage were assessed using a body composition monitor (BF511; Omron Healthcare Co., Ltd., Kyoto, Japan). Waist circumferences (WC) were taken using a nonelastic tape (KaWe Kirchner & Wilhelm GmbH, Asperg, Germany; range 0–150 cm, 1 mm of precision). Measurements were taken twice in a row, considering the average as the result. For blood pressure monitoring, an automatic digital monitor was used (OMRON M3-Intellisense).

Venous blood samples were collected at the baseline in evacuated plastic tubes (VACUETTE TUBE; Greiner Bio-One GmbH, Kremsmunster, Austria). Total cholesterol (TC), LDL-C, high-density lipoprotein cholesterol, triglycerides, apolipoprotein A1, apolipoprotein B (APOB), hs-CRP, HbA1c, inflammatory markers (FN-**γ**, TNF-**α**, IL-10, IL-12, IL-1, IL-6, IL-15), and markers associated with mucosal integrity (l-FAB, calprotectin, and lactoferrin and citrulline, lactulose, mannitol, D-lactate) were measured according to a standardized protocol.

### Microbial profiling

Stool samples were collected and frozen at −80 °C for analysis. DNA was extracted using the QIAamp DNA stool Minikit according to the manufacturer’s instructions (QIAGEN, Hilden, Germany). Microbial analysis of the samples was performed by whole-genome shotgun sequencing in NextSeq Illumina platform, by using a Mid Output kit 2×150pb, and a coverage of ∼ 24 million reads per sample.

The metagenomic analysis was performed following the general guidelines and relying on the bioBakery computational environment. The taxonomic profiling and quantification of organisms’ relative abundances of all metagenomic samples have been quantified using MetaPhlAn 3.062. Metagenomes were mapped internally in MetaPhlAn 3.0 against the marker genes database with BowTie2 version 2.3.4.3 with the parameter “very-sensitive”. The resulting alignments were filtered to remove reads aligned with a MAPQ value <5, representing an estimated probability of the likelihood of the alignments. Functional potential analysis of the metagenomic samples was performed using HUMAnN2 (version 0.11.2 and UniRef database release 2014–07) that computed pathway profiles and gene-family abundances.

For estimating the microbiome species richness of an individual from the taxonomic profiles, we computed three alpha diversity measures: the number of species found in the microbiome (“observed richness”), the Shannon entropy estimation, and the Simpson dominance index using phyloseq (McMurdie & Holmes, 2013). Microbiome dissimilarity between participants (beta diversity) was computed using the weighted UNIFRAC on microbiome taxonomic profiles (Lozupone et al., 2006).

### Microbial co-occurrence networks

Microbial co-occurrence networks were constructed using Sparse InversE Covariance estimation for Ecological Association and Statistical Inference (SpiecEasi) method (Kurtz et al., 2015) with netComi package (Peschel et al., 2021). Nodes represented the microbial species and edge the co-occurrence. Edges connecting nodes were selected by Student’s t-test, and only edges with a significance level after multiple testing adjustments greater than 0.05 were selected. Keystone taxa were defined as highly connected taxa that significantly influenced microbiome structure and function, regardless of their abundance (Banerjee et al., 2018) and were determined as hubs with a degree and betweenness greater than the 90 quartiles.

#### Network comparison

Non-parametric standard permutation tests were used to assess differences between centrality measures and global network characteristics between clusters. A differential association network was constructed using Pearson correlation and Fisher Z test with FDR adjusted P values were used to include only the differentially associated species between clusters.

### Sample Preparation for Metabolomics and UPLC-Q/TOF-MS/MS Procedure

Stools samples were stored in aliquots at −80°C and thawed on ice prior to analysis. The weighted and dry samples were mixed with methanol 80% (1:10 p/v) and mixed for 5 min, then ultrasound was applied for 30 min at room temperature. Finally, samples were centrifugated at 14,800 rpm for 30 min at 4° C, and the supernatant was collected for subsequent detection with Q-TOF in positive mode. Another fraction was evaporated and subsequently resuspended with 40uL of an aqueous solution of acetonitrile (80:20, v/v) for detection with Q-TOF in negative mode.

The UPLC-QTOF-MS analysis was performed by Agilent 1290 Infinity UHPLC with an Agilent 6540 UHD exact mass spectrometer with quadrupole analyzer time of flight (Q/TOF) equipped with an ESI Jet Stream interface.

#### Q-TOF detection in positive mode

Agilent Zorbax Eclipse Plus C18 (1,8µm, 2,1 × 100mm) with an Agilent Zorbax C18 (1,8µm 2.1 × 5 mm) as pre-column was used for chromatographic separation. The column temperature was set at 40°C. The mobile phase consisted of eluent A (0.1% formic acid in aqueous solution) and eluent B (0.1% formic acid in acetonitrile solution). The gradient elution program was set up as follows: 0∼7 min, 30% B; 7∼9 min, 30%→80% B; 9∼11 min, 80%→100%; 11∼13 min, 100% B; 14 min, 100% B with the flow rate at 0.5 mL/min. Detection by TOF MS and Q-TOF MSMS in positive mode. Interval 25-1100 m/z, 5 spectra/s.

#### Q-TOF detection in negative mode

Waters Acquity UPLC BEH Amide column (150 X 2.1 μm; 1.7 μm) with a Waters Acquity UPLC BEH Amide VanGuard Pre-column (5 X 2.1 μm; 1.7 μm) were used for chromatographic separation. The column temperature was set at 45°C. The mobile phase consisted of eluent A (10mM ammonium formate, 0.125% formic acid in water) and eluent B (10mM ammonium formate, 0.125% formic acid in 95:5 acetonitrile:water). The gradient elution program was set up as follows: 0∼2 min, 100% B; 2∼7.7 min, 100%→70%; 7.7∼9.5 min, 70%→40%; 9.5∼10.25 min, 40%→30%; 10.25∼12.75 min, 30%→100%; 12.75∼16.75 min, 100%. Detection by TOF MS in negative mode. Interval 50-1700 m/z, 2 spectra/s, and Q-TOF MSMS in negative mode. Interval 50-1700 m/z, 6 spectra/s.

### Host-microbiome metabolome analysis

Host–microbiome analysis utilized the online tool MetOrigin (*MetOrigin: Discriminating the origins of microbial metabolites for integrative analysis of the gut microbiome and metabolome - Yu - 2022 - iMeta - Wiley Online Library*, s. f.), which integrates gut microbiome data and identifies host, microbiome, and co-metabolism activities associated with microbial metabolites. Microbial species from shotgun metagenomics and metabolomics data were filtered based on a low RSD threshold (30%) and normalized using bacterial relative abundance (percentage) and log-transformed metabolites. The KEGG database was used for the human metabolic pathway curation and a hypergeometric test calculated p-values. Metabolic pathways with log2 p-values > 1 were considered statistically significant and subjected to correlation analysis (Spearman correlation method, p-Value < 0.05). MetOrigin employed Sankey network diagrams to demonstrate associations between microorganisms and metabolites. The analysis generated microbial and metabolite interaction networks by integrating differential metabolites from the host, microbiota, and co-metabolic sources, along with their related bacteria.

### Statistics

All statistical analyses were performed using R version 4.1 (R Core Team, 2022).

#### Multiple-factor analysis

MFA was performed using FactoMineR (Lê et al., 2008) and factoextra R (Kassambara & Mundt, 2020) packages, over the symptomatology, dietary intake, GFD compliance, anthropometric and metabolic measurements to select the most informative variables. Afterwards, a Gaussian mixture model for clustering over the Canberra distance of the selected variables was performed by using mclust R package (Scrucca et al., 2016).

#### Between-group comparisons

Independent two-sample t-test or Wilcoxon rank-sum exact test (in cases where residuals were not reasonably normally distributed) was performed to compare variables between patient groups.

#### Correlation analysis

Pearson’s correlation was run to assess associations between changes in GI symptoms and changes in the gut microbiome (keystone and differential associated taxa), microbial functional potential (differential abundance metaCyc pathways, and GO modules), faecal metabolome (differential abundant metabolites).

#### Statistical significance

For statistical testing, adjusted *P* values below 0.05 were considered statistically significant.

## Results

The baseline characteristics of the 39 participants of the study are displayed in Table 1. Briefly, the majority of participants were female (87%), with an average age of approximately 35 years. All participants exhibited a BMI indicating normal weight. Furthermore, all participants have a confirmed medical diagnosis of celiac disease by biopsy (87%), antibodies (82%) and/or HLA study (45%). The mean time since the diagnosis was 7.1 years, and the mean time undergoing a GFD was 7.2 years (range 1 - 28 years). Moreover, all participants reported experiencing gastrointestinal or extraintestinal symptoms characteristic of CD, with an average symptom count of 3 different symptoms (range 1 - 8 different symptoms).

**Table 1.** Baseline characteristics from patients included in the study.

| **Characteristic** |  | **N =39<sup>a</sup>** |
| --- | --- | --- |
| Male |  | 5 (13%) |
| Female |  | 34 (87%) |
| Age |  | 35 (11) |
| Weight |  | 61 (12) |
| Height |  | 164 (7) |
| BMI |  | 22.6 (4.2) |
| Years since diagnosis |  | 10 (7) |
| Positive biopsy |  | 34 (87%) |
| Positive Antibodies |  | 32 (82%) |
| Positive HLA study |  | 17 (45%) |
| Years under GFD |  | 10 (7) |
| Number of<br>different<br>symptoms | 1 | 6 (15%) |
|  | 2 | 9 (23%) |
|  | 3 | 6 (15%) |
|  | 4 | 5 (13%) |
|  | 5 | 7 (18%) |
|  | >6 | 5 (16%) |

<sup>a</sup>Mean (SD). N(%). BMI, Body mass index. HLA, Human leukocyte antigens. GFD, Gluten free diet.

### NRCD subgroups can be defined according to clinical and demographic variables not influenCD by gluten consumption

Multiple Factorial Analysis was used to determine the importance of the clinical, demographic, inflammatory, and mucosal integrity biomarkers. From the initial set of 99 variables analysed in the study, 47 were chosen based on their informativeness (data not shown). These selected variables were subjected to a Gaussian mixture model analysis, utilizing the Canberra distance metric, leading to the identification of two distinct clusters.

Cluster 1, designated as “Low-grade symptoms,” exhibited lower levels of symptoms and inflammatory markers. Conversely, Cluster 2, termed “High-grade symptoms,” displayed elevated scores in patient-reported symptom questionnaires, heightened inflammatory markers, and increased intestinal permeability. A comprehensive summary of these findings can be found in Table 2.

**Table 2.**
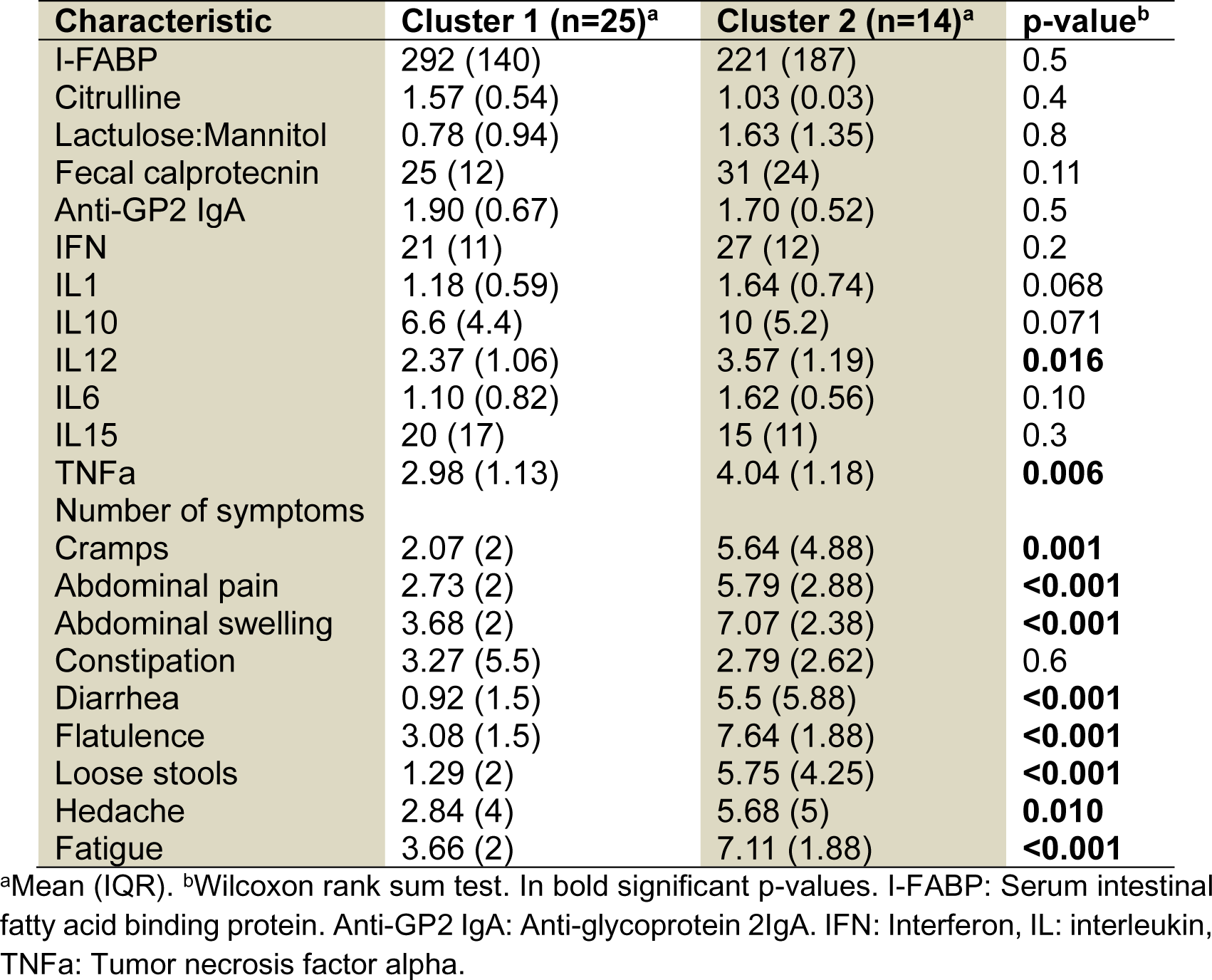
Barrier integrity, inflammatory markers and symptoms from patients allocated in each cluster.

| Characteristic | Cluster 1 (n=25)<sup>a</sup> | Cluster 2 (n=14)<sup>a</sup> | p-value<sup>b</sup> |
| --- | --- | --- | --- |
| I-FABP | 292 (140) | 221 (187) | 0.5 |
| Citrulline | 1.57 (0.54) | 1.03 (0.03) | 0.4 |
| Lactulose:Mannitol | 0.78 (0.94) | 1.63 (1.35) | 0.8 |
| Fecal calprotectin | 25 (12) | 31 (24) | 0.11 |
| Anti-GP2 IgA | 1.90 (0.67) | 1.70 (0.52) | 0.5 |
| IFN | 21 (11) | 27 (12) | 0.2 |
| IL1 | 1.18 (0.59) | 1.64 (0.74) | 0.068 |
| IL10 | 6.6 (4.4) | 10 (5.2) | 0.071 |
| IL12 | 2.37 (1.06) | 3.57 (1.19) | **0.016** |
| IL6 | 1.10 (0.82) | 1.62 (0.56) | 0.10 |
| IL15 | 20 (17) | 15 (11) | 0.3 |
| TNFα | 2.98 (1.13) | 4.04 (1.18) | **0.006** |
| Number of symptoms |  |  |  |
| Cramps | 2.07 (2) | 5.64 (4.88) | **0.001** |
| Abdominal pain | 2.73 (2) | 5.79 (2.88) | **<0.001** |
| Abdominal swelling | 3.68 (2) | 7.07 (2.38) | **<0.001** |
| Constipation | 3.27 (5.5) | 2.79 (2.62) | 0.6 |
| Diarrhea | 0.92 (1.5) | 5.5 (5.88) | **<0.001** |
| Flatulence | 3.08 (1.5) | 7.64 (1.88) | **<0.001** |
| Loose stools | 1.29 (2) | 5.75 (4.25) | **<0.001** |
| Hedache | 2.84 (4) | 5.68 (5) | **0.010** |
| Fatigue | 3.66 (2) | 7.11 (1.88) | **<0.001** |

<sup>a</sup>Mean (IQR). <sup>b</sup>Wilcoxon rank sum test. In bold significant p-values. I-FABP: Serum intestinal fatty acid binding protein. Anti-GP2 IgA: Anti-glycoprotein 2IgA. IFN: Interferon, IL: interleukin, TNFα : Tumor necrosis factor alpha.

An in-depth analysis of the symptomatology revealed noteworthy distinctions between the two clusters. Patients in cluster 1 experienced an average of two different symptoms, while patients in cluster 2 reported up to five different symptoms (*P* < 0.01). Particularly, individuals assigned to cluster 2 displayed significantly more severe gastrointestinal symptoms, including cramps, abdominal pain, abdominal swelling, constipation, diarrhoea, flatulence, and loose stools, as assessed by two distinct questionnaires (*P* < 0.001). Furthermore, patients in cluster 2 reported a higher incidence of extraintestinal symptoms such as headaches and fatigue (*P* < 0.001).

To further investigate the integrity of the gastrointestinal barrier, we conducted relevant measurements. Although a noticeable trend suggesting greater barrier damage among participants in cluster 2 was observed, the results did not reach statistical significance.

In terms of inflammatory biomarkers, our analysis revealed elevated levels of IL-12 and TNFa in patients belonging to cluster 2 (*P* < 0.001). These findings indicate a potential association between cluster 2 and heightened inflammatory activity, warranting further investigation.

Lastly, it is important to note that no significant differences were observed in the CDAT questionnaire (mean punctuation 11.8 points; 13.5 points, *P*=ns) or the faecal gluten immunogenic peptides (0.53; 0.10, *P*=ns), indicating low overall gluten consumption among the study participants.

### Diet quality was fair in both clusters of patients

Micro and macronutrient consumption of each participant was calculated by using self-report questionnaires supervised by a nutritionist. The diet quality of patients in both clusters was fair, as evidenced by a Healthy Eating Index close to 50% (Cluster 1: 55%, Cluster 2: 53%; *P*=ns). Deficiencies primarily arise from low cereal consumption and limited dietary variety (Table 3). Regarding macronutrient intake, both groups display similar consumption levels of fats, carbohydrates, and simple sugars. However, the percentage of protein relative to the total caloric value of the diet was lower in patients from Cluster 2 compared to Cluster 1.

**Table 3.** Nutritional characteristic of each cluster.

| Characteristic | Cluster 1 (n=25)<sup>a</sup> | Cluster 2 (n=14)<sup>a</sup> | p-value<sup>b</sup> |
| --- | --- | --- | --- |
| CDAT |  |  |  |
| HEI | 55 (10) | 53 (13) | 0.4 |
| Diet variety |  |  |  |
| Calories | 2,094 (532) | 2,201 (519) | 0.6 |
| Proteins | 82 (20) | 78 (21) | 0.5 |
| Carbohydrates | 184 (64) | 194 (49) | 0.7 |
| % Sugars |  |  |  |
| % Fat |  |  |  |
| Cereals |  |  |  |
| Soluble fibre | 2.72 (1.34) | 3.29 (2.30) | 0.6 |
| Insoluble fibre | 5.57 (2.33) | 6.53 (3.77) | 0.4 |
| Vegetal fibre | 22 (6) | 22 (10) | 0.7 |
| Zinc |  |  |  |
| Calcium | 717 (268) | 678 (230) | 0.7 |
| Iron | 10.7 (3.5) | 9.7 (3.3) | 0.7 |
| Magnesium | 237 (64) | 247 (90) | 0.9 |
| Vitamin K | 144 (84) | 126 (70) | 0.7 |
| Vitamin D | 3.7 (3.7) | 2 (2.7) | **0.022** |
| Vitamin B12 | 6.89 (10.08) | 4.16 (2.236) | 0.3 |
| Vitamin A | 1,305 (2,506) | 815 (452) | 0.7 |
| Vitamin E | 10.3 (4.1) | 13.8 (8.7) | 0.4 |
| Folic Acid | 226 (93) | 203 (94) | 0.6 |
| Quercetin |  |  |  |

<sup>a</sup>Mean (IQR). <sup>b</sup>Wilcoxon rank sum test. In bold significant p-values. CDAT: Celiac Dietary Adherence Test. HEI: Healthy eating index.

Both groups demonstrate adequate intake percentages (above 100%) for major vitamins. However, it is noteworthy that vitamin D consumption was low (74% and 39% adequacy relative to the RDI for each cluster; P < 0.05). The adequacy percentage for zinc, iron, and calcium was around 60% of the RDI, with no significant differences between groups. Regarding other nutrients, we only observed significant differences in quercetin consumption between both groups.

### Microbial community structure is different between clusters

The stool samples from the patients underwent shotgun metagenomics analysis to unravel the microbial composition. Following a thorough analysis, a total of 7,685,850 high-quality sequences were identified, unveiling the presence of 308 distinct microbial species. The relative abundance analysis revealed the dominance of the phyla *Bacteroidetes*, *Firmicutes*, *Verrucomcirobia*, *Proteobacteria* and *Actinobacteria* within the microbial community. Furthermore, we computed alpha diversity indexes and observed no discernible differences across the clusters, indicating comparable microbial diversity (Chao index 186 vs. 202; Simpson 0.92 vs. 0.93; Shannon 3.15 vs 3.33 *P*=ns).

To comprehensively explore the microbial community structure, sparse inverse covariance estimation and model selection techniques were employed to construct co-occurrence networks. Intriguingly, distinct network properties were observed for each cluster, highlighting the presence of keystone taxa that exerted significant influence over the community dynamics (Figure 1). Furthermore, we constructed a differential association network, linking nodes based on their differential associations between the two clusters, as determined through Pearson correlation analysis. To specifically focus on the associations of interest, we constructed association networks solely comprising species that exhibited differential associations (Figure 2).

**Figure 1:**
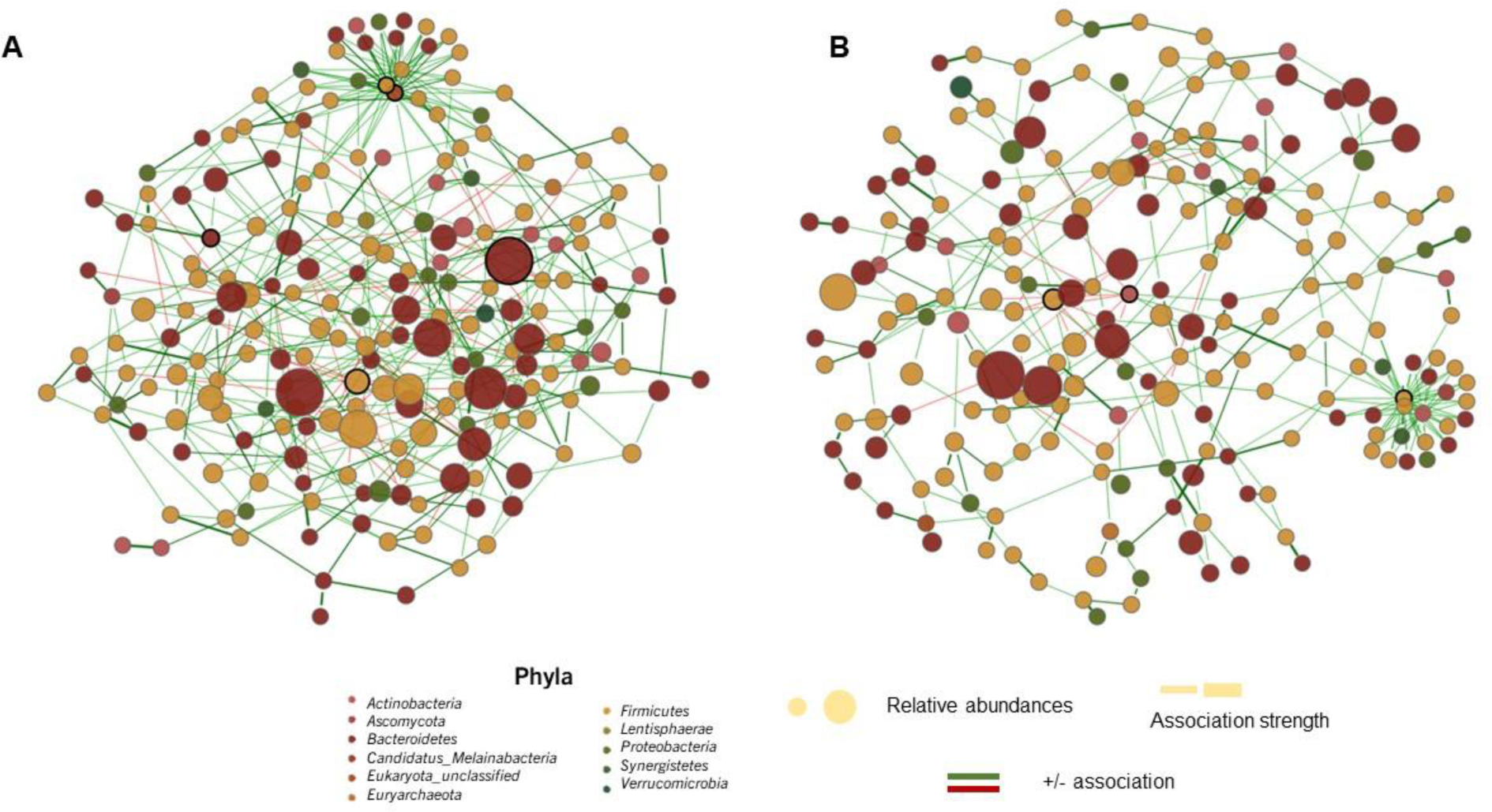
Co-occurrence networks of the microbiome of patients of low-grade symptoms (A) and high-grade symptoms (B) clusters. Co-occurrence network of each cluster. The nodes in the graph represent different bacterial species. The size of each node corresponds to its relative abundance, while the edges represent statistically significant associations between the nodes (P < 0.05). Green edges indicate positive relationships, while red edges indicate negative ones. The thickness of each edge indicates the strength of the association. Nodes highlighted in figure a correspond to keystone taxa calculated as nodes with betweenness and degree greater than Quantile 0.90

**Figure 2.**
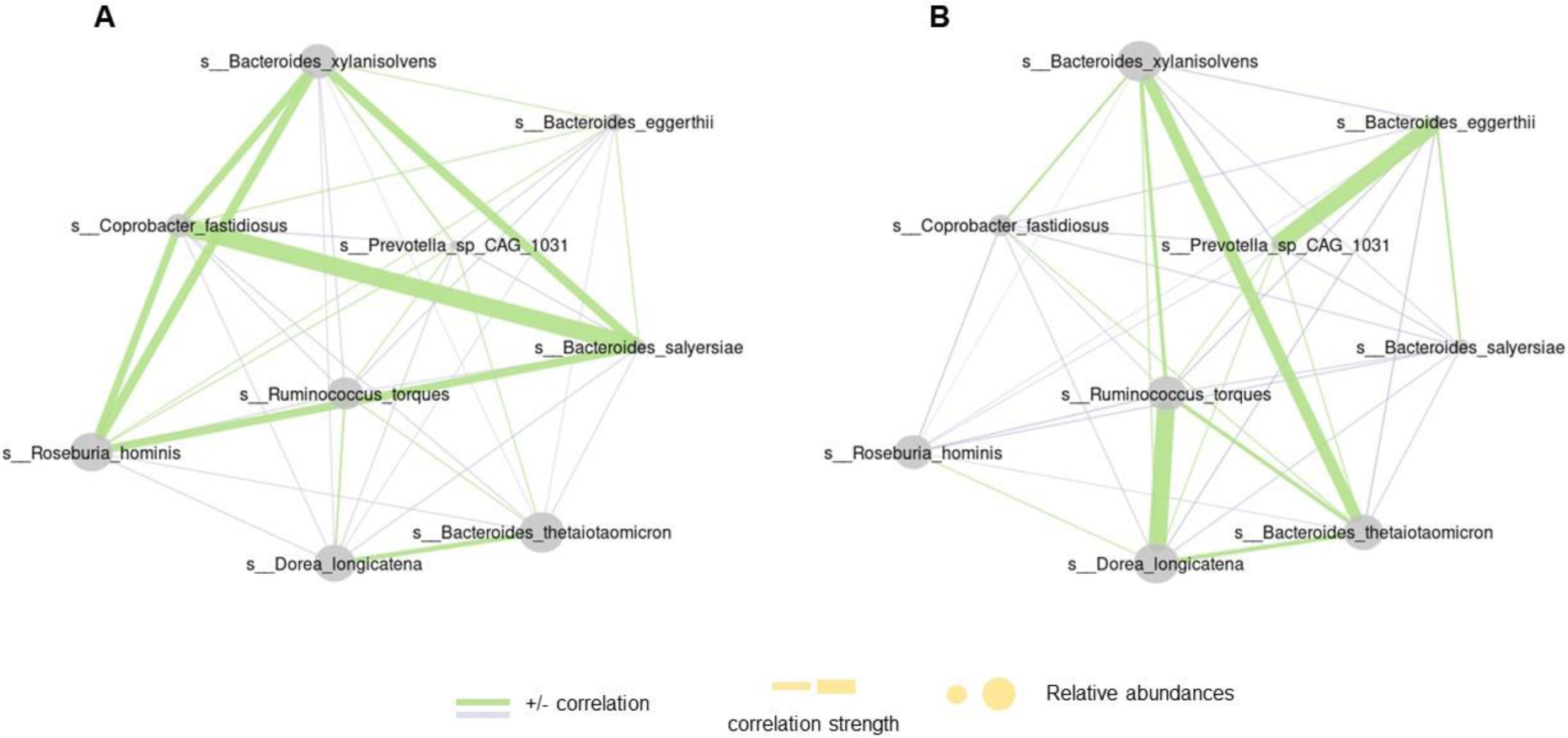
Network of different associated species in of low-grade symptoms (A) and high-grade symptoms (B) clusters.

By integrating these analytical approaches, we gained a greater understanding of the microbial community structure and the intricate associations among different taxa. This comprehensive exploration enhances our insights into the functional dynamics and ecological interactions within the microbial ecosystem of the two patient clusters.

### Metabolome analysis and functional microbiome profiling

A total of 372 metabolites were identified by HPLC and were classified into four groups: 5 host (human)-specific metabolites, 56 microbial metabolites, 103 microbial-host cometabolites, and 208 others (57 drug, 48 food, 1 environmental related and 102 unknown). There were 36 differential metabolites between the two clusters, including 6 microbial metabolites, 12 host-microbial cometabolites, and 18 others. A metabolic pathway enrichment analysis (MPEA) was performed with the microbial and host-microbial cometabolites (Figure 3). There were 4, and 29 related metabolic pathways matched against microbial and host-microbial metabolites pathway databases, respectively. Among these, lysine biosynthesis (microbial) and Glutathione metabolism, Glyoxylate and dicarboxylate metabolism, C5-Branched dibasic acid metabolism, Alanine, aspartate and glutamate metabolism, Pentose phosphate pathway, Butanoate metabolism, D-Amino acid metabolism, Arginine and proline metabolism, and Caffeine metabolism (Co-metabolism) were significantly different between clusters (*P* <0.05).

**Figure 3.**
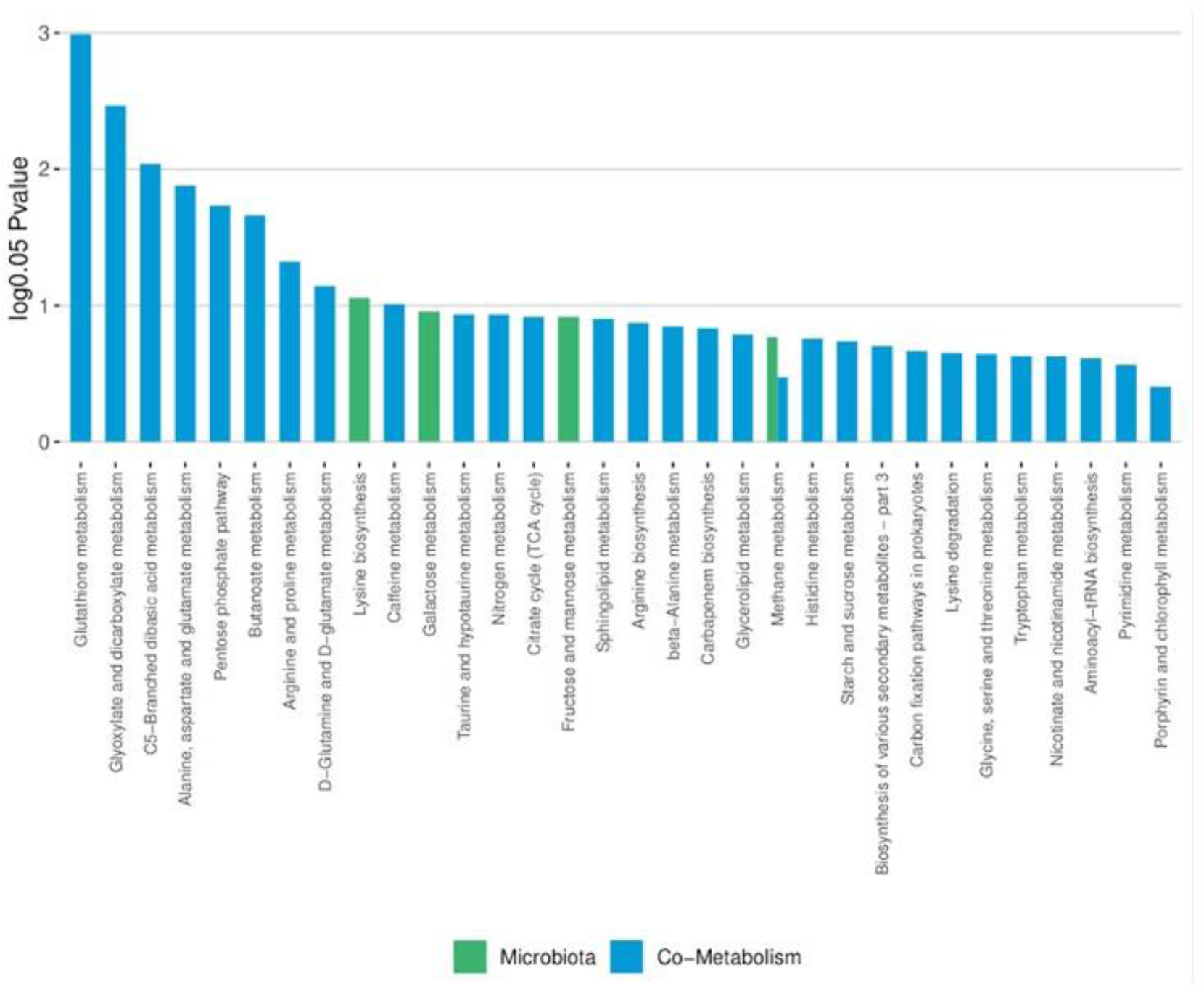
Metabolite pathway enrichment analysis of microbiota-derived and co-metabolism erived metabolites.

Through Sankey network analysis, we were able to link all the possible bacteria that may participate in a specific metabolic reaction, allowing us to identify the interplay between bacteria and metabolites. Finally, we summarised and integrated differential metabolites from microbiota and cometabolism origins and their related bacteria within a network to obtain a whole picture of the microbiome and metabolome cross-talk highlighting the interactions with both biological and statistical signification (Figure 4). We found that *Aanesostipes hadrus* and *Lawsonibacter asaccharolyticus* were associated with the regulation of several metabolites implied in the abovementioned metabolic pathways.

**Figure 4:**
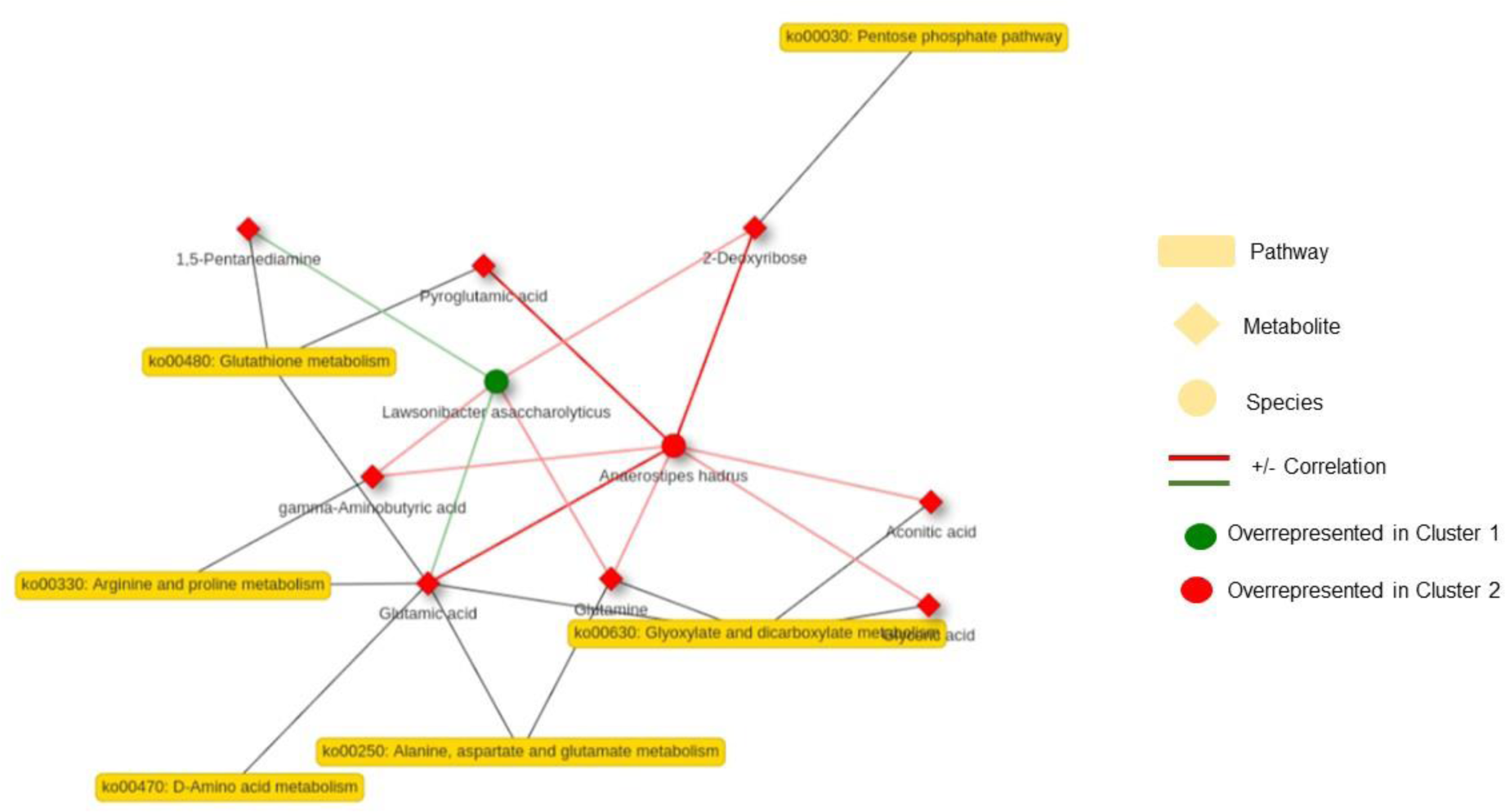
Network integrating differential metabolites from microbiota and cometabolism origins and their related bacteria.

### Correlations between gastrointestinal symptoms and faecal microbiome and metabolome reveal an association between symptom severity and microbial-derived metabolites

The potential association between changes in symptom scores and changes in gut microbiome (keystone and differential associated taxa), microbial functional potential (differential abundance metaCyc pathways, and GO modules), and faecal metabolome (differential abundant metabolites) were assessed by performing Pearson’s correlation test (Figure 5).

**Figure 5:**
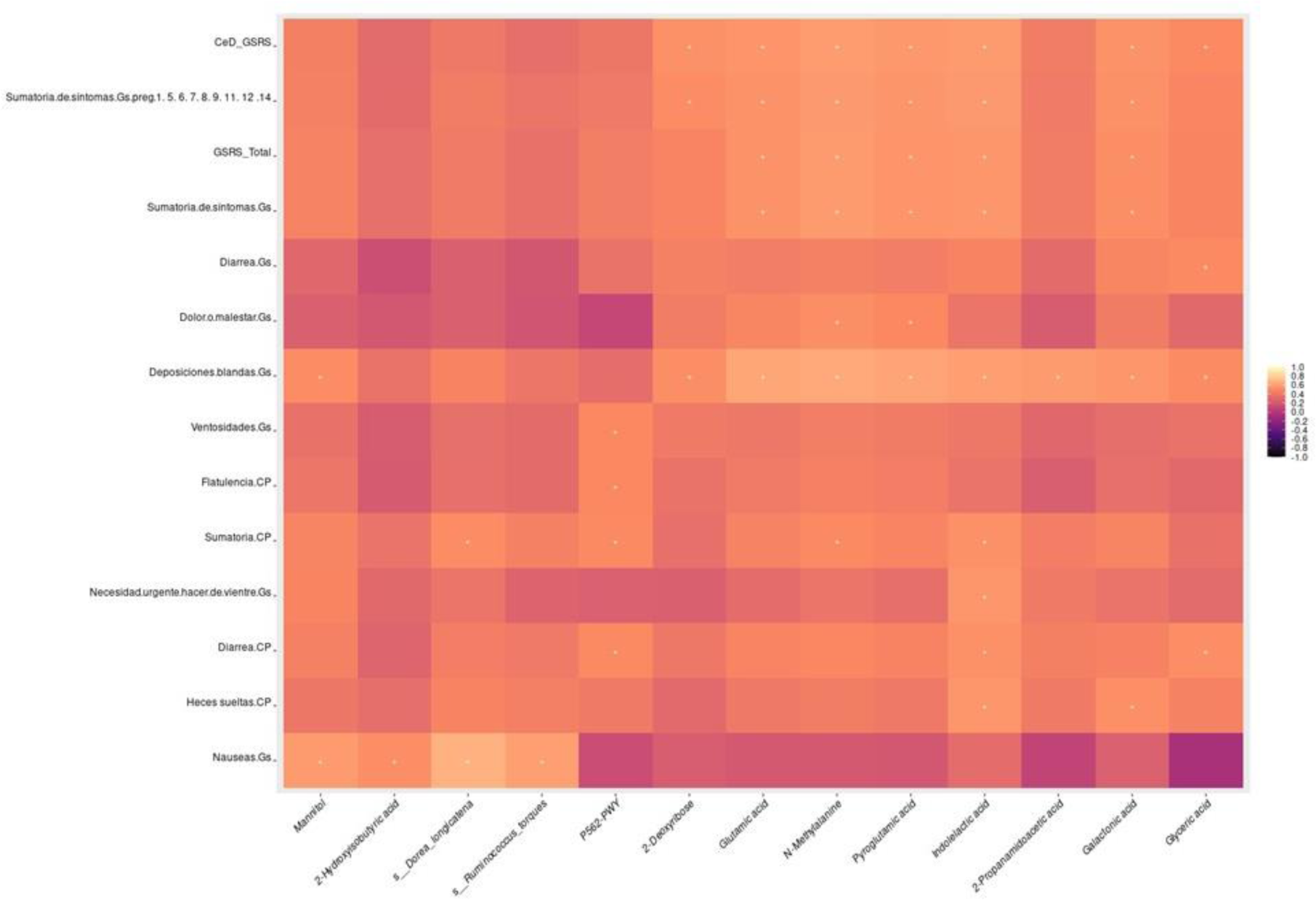
Pearson’s correlation test between changes in symptom scores and changes in gut microbiome (keystone and differential associated taxa), microbial functional potential (differential abundance metaCyc pathways, and GO modules), and faecal metabolome (differential abundant metabolites)

We identified a strong correlation between the punctuation and total score of the GSRS questionnaire (both CD-related symptoms and all gastrointestinal symptoms) with specific faecal metabolites, namely glutamic acid, N-methylamine, pyroglutamic acid, indolelactic acid, and galactonic acid (*P* <0.05). These metabolites were found to be more abundant in the faecal metabolome of patients belonging to Cluster 2. On the other hand, the overall punctuation of the CD-Pro symptom questionnaire demonstrated associations with *Dorea longicatena*, myo-inositol degradation pathway, N-methylamine and indolelactic acid.

Moreover, certain associations with symptoms were observed. In particular, general pain showed associations with N-methylamine and pyroglutamic acid. Loose stools were associated with indolelactic and galactonic acid. Flatulence and diarrhoea were associated with the Myo-inositol degradation pathway. Additionally, nauseous symptoms were linked to Mannitol, 2-hydroxybutyric acid, *Dorea longicatena*, and *Ruminococcus torques* (*P* <0.05). These findings indicate a potential relationship between the faecal microbiome and metabolome and symptomatology, shedding light on the significance of specific metabolites in the context of gastrointestinal health and their possible role in symptom manifestation.

## Discussion

The gut microbiome has garnered significant attention in the context of active CD, where its pivotal role, particularly a dysbiotic state marked by an abundance of gram-negative bacteria like *Bacteroidetes* and *Proteobacteria*, has been extensively explored (Arcila-Galvis et al., 2022). However, in the case of NRCD, the scientific understanding remains relatively limited. Nevertheless, emerging evidence hints at the potential influence of Small Intestinal Bacterial Overgrowth (SIBO) on the persistence of symptoms in these patients (Leffler et al., 2007; O’Mahony et al., 1996; Penny et al., 2020). While this area requires further investigation, it underscores the relevance of exploring the gut microbiome’s involvement in the persistence of symptoms in CD patients despite maintaining a GFD.

This study presents novel insights into NRCD disease patients, highlighting the presence of distinct clusters characterized by variations in symptoms, inflammatory markers, microbial community structure, and associations between microbial metabolites and symptoms.

Our analysis revealed no significant differences in terms of mucosal integrity between the two groups under investigation. Both groups exhibited evidence of impaired permeability, as indicated by a high Lactulose:Mannitol ratio (0.78 vs 1.63, *P*=ns), representing an approximately 20-fold increase compared to values observed in healthy controls (Benjamin et al., 2008). Additionally, enterocyte damage was apparent in both groups, as reflected by low levels of citrulline (1.57 vs 1.03 /dL, *P*=ns), which has been associated with Advanced degrees of villous atrophy. However, it is noteworthy that both groups displayed levels of faecal calprotectin (25, 31 *P*=ns) and Intestinal Fatty Acid Binding Protein (292,221 *P*=ns) within the normal range, suggesting that despite the mucosal damage, the enterocytes maintain their functionality. These findings indicate that factors other than enterocyte functionality contribute to the persistence of symptoms, affecting membrane permeability and inflammatory processes.

We then further investigate the microbial composition of the two clusters. The use of microbial co-occurrence networks provides a comprehensive approach to investigating the structure of microbial communities, surpassing the limitations of differential abundance analyses. This network-based analysis enables the identification of keystone taxa, which play crucial roles within the community and may otherwise remain unnoticed using alternative methods (Banerjee et al., 2018). In this study, despite the comparable alpha diversity values observed among the microbial communities under investigation, the exploration of community structure through network analysis has unveiled two distinct communities harbouring specific microbial entities that could potentially impact symptom persistence. While subtle differences in microbial composition were observed, network-based characterization has offered valuable insights into the intricate dynamics and potential functional implications of these microbial communities. Such findings highlight the significance of network-based approaches in unravelling the complex interplay between microbial taxa and their potential influence on disease outcomes.

Notably, we elucidate the characteristics of a “Low-grade symptoms cluster” where specific microorganisms may play a role in the regulation of the immune response through diverse mechanisms. *Blastocystis* sp., subtype 1, exhibits the ability to modulate pro and anti-inflammatory cytokines via exosomes (Norouzi et al., 2022). *Dialister* sp. CAG357 and *Ruminococcus bromii* contribute through butyrate production (Ze et al., 2012), while *Bacteroides* sp. OM08-11 and *B. vulgatus* decrease lipopolysaccharide production, which is implicated in TLR-4 activation. Furthermore, *Roseburia hominis* strengthens gut barrier function (Patterson et al., 2017). These findings suggest the presence of a microbial community associated with the homeostasis of the immune response and the maintaining of the barrier integrity lowering the severity of inflammation and gastrointestinal symptoms.

Conversely, the “High-grade symptoms cluster” displays a distinct gut microbiome composition characterized by lactate-producing microorganisms (*Coprobacillus cateniformis* and *Fusicatenibacter saccharivorans*), *Eggerthella lenta* associated with intestinal Th17 activation (Alexander et al., 2022), and *Ruminococcus torques* previously linked to compromised gut barrier integrity (Rajilic-Stojanovic & de Vos, 2014). Th17 immune response, known to play a pivotal role in autoimmune and inflammatory diseases, including CD pathogenesis, is marked by increased expression of Th17-related cytokines in active CD patients (Castellanos-Rubio et al., 2009). Recent findings of gluten-specific IL-17A-producing cells in the duodenum of CD patients further underscore its relevance. *E. lenta* and lactate, both present in cluster 2 patients, have been associated with Th17 activation (Alexander et al., 2022). Additionally, *R. torques*, a potent mucus degrader, has been linked to the severity of irritable bowel symptoms. Notably, patients in cluster 2 exhibit elevated levels of pro-inflammatory cytokines, specifically IL-12 and TNFa, these findings, alongside the microbial community structure, suggest a microbiome-mediated pro-inflammatory state independent of gluten consumption.

Importantly, it is worth noting that a general deficiency in vitamin D was observed, with cluster 2 patients displaying more pronounced insufficiency. Vitamin D, a crucial micronutrient in immune response maintenance, has been associated with celiac disease and plays a role in shaping the composition and structure of the gut microbiome.

Previous research has established that CD is characterized by metabolic alterations encompassing impaired energy metabolism, malabsorption, and perturbations in the gut microbiota (Ryan et al., 2015). In our metabolomic analysis, we aimed to distinguish between microbial and host-derived metabolites, as well as co-metabolites resulting from their interactions. Notably, patients belonging to cluster two exhibited elevated levels of metabolites associated with tryptophan metabolism (Indoxyl sulfate, Indolelactic acid), protein hydrolysis (cadaverine), bacterial cell wall components (2,6-Diaminopimelic acid), and inflammation (mannitol). Through metOrigin analysis (*MetOrigin: Discriminating the origins of microbial metabolites for integrative analysis of the gut microbiome and metabolome - Yu - 2022 – iMeta - Wiley Online Library*, s. f.), we gained insight into the intricate interplay between the metabolome and microbiome by integrating the metabolic pathways of the microbiome with the detected metabolites in the samples. Our findings revealed that pathways mainly related to amino acid metabolism were enriched, and some microbial species (*A. hadrus* and *L. asaccharolyticus*) were significantly related to the metabolites participating in those pathways. Abnormalities in amino acid metabolism has been associated with impairment in the gut barrier integrity through activation of the immune response by Th17 activation (Castellanos-Rubio et al., 2009).

Furthermore, we conducted a comprehensive investigation into the association between microbial taxa, differential metabolites, metabolic pathways, and the observed symptoms. Our analysis revealed a strong association between microbial-derived metabolites (mannitol, indolelactic acid and galactonic acid), human-microbial co-metabolism-derived metabolites (2-Deoxyribose, glutamic acid, pyroglutamic acid and Glyceric acid) and gastrointestinal symptoms, showing that higher levels of these metabolites are associated with higher scores in partial and total questionnaires. We also found that the Myo-inositol degradation pathway was significantly abundant in patients from cluster two (*P*<0.005) and was associated with the presence of flatulence, diarrhoea and the summary of symptoms of the questionnaires. Myo-inositol degradation is essential for carbon assimilation in bacteria, and it serves as a key player in various metabolic functions. Inadequate myo-inositol levels have been linked to heightened inflammation and apoptosis in the intestinal mucosa, along with reduced cell proliferation and antioxidant capacity. This pathway exhibits activity exclusively in specific kidney cell types and assumes significance, particularly under certain disease conditions. Here we show for the first time its possible association with CD symptoms.

Association between symptoms and particular microbial species was only found for *Dorea loncitanea* and *Ruminoccous torques* and the presence of nausea. These microbial species were more abundant in the gut microbiome of patients from cluster two and were also differentially associated in the microbial network (Figure 2), showing a strong positive correlation and suggesting a possible upregulation.

Prior research has established a correlation between alterations in the gut microbiome composition and the occurrence or intensification of various gastrointestinal symptoms (REF). Our findings support the hypothesis that the metabolic capabilities of the microbiome, specifically reflecting the functional aspects of the microbial community, exert a significant influence on the persistence of symptoms. This influence is primarily mediated through the regulation of amino acid metabolism and immune homeostasis. Of particular importance, we underscore the involvement of specific microorganisms and their associated metabolites in triggering immune activation, which subsequently leads to alterations in the epithelial barrier. These changes in the barrier function may give rise to visceral hypersensitivity and abnormalities in gut motility, thereby manifesting as severe symptomatology.

In conclusion, this study contributes to the understanding of NRCD by highlighting the role of the gut microbiome, mucosal integrity, and metabolomic profiles in symptom persistence. Further research in this field will deepen our knowledge of the intricate interplay between these factors and may pave the way for targeted interventions to improve clinical outcomes in NRCD patients.

## Declarations

### Ethics approval and consent to participate

The study was approved by the Research Ethics Committee of the IMDEA Food Foundation (PI-032; Approval date: June 12th, 2017). All the participants have signed the informed consent and donated the samples for future research in the same line of research. These samples are a collection: Reference: C.0004841. Registro Nacional de Biobancos Hospital Carlos III. PI: Dr. Viviana Loria Kohen. Date: 21/03/2019.

## References

Abdulkarim, A. S., Burgart, L. J., See, J., & Murray, J. A. (2002). Etiology of nonresponsive celiac disease: Results of a systematic approach. The American Journal of Gastroenterology, 97(8), 2016–2021. 10.1111/j.1572-0241.2002.05917.x

Alexander, M., Ang, Q. Y., Nayak, R. R., Bustion, A. E., Sandy, M., Zhang, B., Upadhyay, V., Pollard, K. S., Lynch, S. V., & Turnbaugh, P. J. (2022). Human gut bacterial metabolism drives Th17 activation and colitis. Cell Host & Microbe, 30(1), 17–30.e9. 10.1016/j.chom.2021.11.001

Al-Toma, A., Volta, U., Auricchio, R., Castillejo, G., Sanders, D. S., Cellier, C., Mulder, C. J., & Lundin, K. E. A. (2019). European Society for the Study of Coeliac Disease (ESsCD) guideline for coeliac disease and other gluten-related disorders. United European Gastroenterology Journal, 7(5), 583–613. 10.1177/2050640619844125

Arcila-Galvis, J. E., Loria-Kohen, V., Ramírez de Molina, A., Carrillo de Santa Pau, E., & Marcos-Zambrano, L. J. (2022). A comprehensive map of microbial biomarkers along the gastrointestinal tract for celiac disease patients. Frontiers in Microbiology, 13, 956119. 10.3389/fmicb.2022.956119

Banerjee, S., Schlaeppi, K., & van der Heijden, M. G. A. (2018). Keystone taxa as drivers of microbiome structure and functioning. Nature Reviews. Microbiology, 16(9), 567–576. 10.1038/s41579-018-0024-1

Bascuñán, K. A., Araya, M., Roncoroni, L., Doneda, L., & Elli, L. (2020). Dietary Gluten as a Conditioning Factor of the Gut Microbiota in Celiac Disease. Advances in Nutrition (Bethesda, Md.), 11(1), 160–174. 10.1093/advances/nmz080

Benjamin, J., Makharia, G. K., Ahuja, V., Kalaivani, M., & Joshi, Y. K. (2008). Intestinal permeability and its association with the patient and disease characteristics in Crohn’s disease. World Journal of Gastroenterology, 14(9), 1399–1405. 10.3748/wjg.14.1399

Caio, G., Volta, U., Sapone, A., Leffler, D. A., De Giorgio, R., Catassi, C., & Fasano, A. (2019). Celiac disease: A comprehensive current review. BMC Med., 17(1), 142.

Castellanos-Rubio, A., Santin, I., Irastorza, I., Castaño, L., Carlos Vitoria, J., & Ramon Bilbao, J. (2009). TH17 (and TH1) signatures of intestinal biopsies of CD patients in response to gliadin. Autoimmunity, 42(1), 69–73. 10.1080/08916930802350789

Catassi, C., Verdu, E. F., Bai, J. C., & Lionetti, E. (2022). Coeliac disease. Lancet (London, England), 399(10344), 2413–2426. 10.1016/S0140-6736(22)00794-2

Dahl, W. J., Rivero Mendoza, D., & Lambert, J. M. (2020). Diet, nutrients and the microbiome. Progress in molecular biology and translational science, 171, 237–263. 10.1016/BS.PMBTS.2020.04.006

Fueyo-Díaz, R., Magallón-Botaya, R., Sánchez-Calavera, M. A., Asensio-Martínez, A., & Gascón-Santos, S. (2015). Protocol development for a scale to assess self-efficacy in adherence to a gluten free diet: Self-Efficacy and Celiac Disease Scale (Vol. 19).

Kassambara, A., & Mundt, F. (2020). factoextra: Extract and Visualize the Results of Multivariate Data Analyses. https://CRAN.R-project.org/package=factoextra

Kurtz, Z. D., Müller, C. L., Miraldi, E. R., Littman, D. R., Blaser, M. J., & Bonneau, R. A. (2015). Sparse and compositionally robust inference of microbial ecological networks. PLoS Computational Biology, 11(5), e1004226. 10.1371/journal.pcbi.1004226

Lê, S., Josse, J., & Husson, F. (2008). FactoMineR: A Package for Multivariate Analysis. Journal of Statistical Software, 25(1), 1–18. 10.18637/jss.v025.i01

Leffler, D. A., Dennis, M., Hyett, B., Kelly, E., Schuppan, D., & Kelly, C. P. (2007). Etiologies and Predictors of Diagnosis in Nonresponsive Celiac Disease. Clinical Gastroenterology and Hepatology, 5(4), 445–450. 10.1016/j.cgh.2006.12.006

Leffler, D. A., Kelly, C. P., Green, P. H. R., Fedorak, R. N., DiMarino, A., Perrow, W., Rasmussen, H., Wang, C., Bercik, P., Bachir, N. M., & Murray, J. A. (2015). Larazotide acetate for persistent symptoms of celiac disease despite a gluten-free diet: A randomized controlled trial. Gastroenterology, 148(7), 1311–9.e6.

Leonard, M. M., Cureton, P., & Fasano, A. (2017). Indications and Use of the Gluten Contamination Elimination Diet for Patients with Non-Responsive Celiac Disease. Nutrients, 9(10).

Levy, M., Blacher, E., & Elinav, E. (2017). Microbiome, metabolites and host immunity. Curr. Opin. Microbiol., 35, 8–15.

Lozupone, C., Hamady, M., & Knight, R. (2006). UniFrac – An online tool for comparing microbial community diversity in a phylogenetic context. BMC Bioinformatics, 7(1), Article 1. 10.1186/1471-2105-7-371

McMurdie, P. J., & Holmes, S. (2013). phyloseq: An R Package for Reproducible Interactive Analysis and Graphics of Microbiome Census Data. PLoS ONE, 8(4), Article 4. 10.1371/journal.pone.0061217

MetOrigin: Discriminating the origins of microbial metabolites for integrative analysis of the gut microbiome and metabolome—Yu—2022—iMeta—Wiley Online Library. (s. f.). Recuperado 25 de julio de 2023, de https://onlinelibrary.wiley.com/doi/full/10.1002/imt2.10

Norouzi, M., Pirestani, M., Arefian, E., Dalimi, A., Sadraei, J., & Mirjalali, H. (2022). Exosomes secreted by Blastocystis subtypes affect the expression of proinflammatory and anti-inflammatory cytokines (TNFα, IL-6, IL-10, IL-4). Frontiers in Medicine, 9, 940332. 10.3389/fmed.2022.940332

O’Mahony, S., Howdle, P. D., & Losowsky, M. S. (1996). Review article: Management of patients with non-responsive coeliac disease. Alimentary Pharmacology & Therapeutics, 10(5), 671–680. 10.1046/j.1365-2036.1996.66237000.x

Ortega, R. M., Pérez-Rodrigo, C., & López-Sobaler, A. M. (2015). Dietary assessment methods: Dietary records. Nutricion Hospitalaria, 31 Suppl 3, 38–45. 10.3305/nh.2015.31.sup3.8749

Patterson, A. M., Mulder, I. E., Travis, A. J., Lan, A., Cerf-Bensussan, N., Gaboriau-Routhiau, V., Garden, K., Logan, E., Delday, M. I., Coutts, A. G. P., Monnais, E., Ferraria, V. C., Inoue, R., Grant, G., & Aminov, R. I. (2017). Human Gut Symbiont Roseburia hominis Promotes and Regulates Innate Immunity. Frontiers in Immunology, 8, 1166. 10.3389/fimmu.2017.01166

Penny, H. A., Baggus, E. M. R., Rej, A., Snowden, J. A., & Sanders, D. S. (2020). Non-Responsive Coeliac Disease: A Comprehensive Review from the NHS England National Centre for Refractory Coeliac Disease. Nutrients, 12(1), 216. 10.3390/nu12010216

Peschel, S., Müller, C. L., von Mutius, E., Boulesteix, A.-L., & Depner, M. (2021). NetCoMi: Network construction and comparison for microbiome data in R. Briefings in Bioinformatics, 22(4). 10.1093/bib/bbaa290

R Core Team. (2022). R: A Language and Environment for Statistical Computing. R Foundation for Statistical Computing. https://www.R-project.org/

Rajilic-Stojanovic, M., & de Vos, W. M. (2014). The first 1000 cultured species of the human gastrointestinal microbiota. FEMS Microbiol. Rev., 38(5), 996–1047.

Ryan, D., Newnham, E. D., Prenzler, P. D., & Gibson, P. R. (2015). Metabolomics as a tool for diagnosis and monitoring in coeliac disease. Metabolomics, 11(4), 980–990. 10.1007/s11306-014-0752-9

Sansotta, N., Amirikian, K., Guandalini, S., & Jericho, H. (2018). Celiac Disease Symptom Resolution: Effectiveness of the Gluten-free Diet. Journal of Pediatric Gastroenterology and Nutrition, 66(1), 48–52. 10.1097/MPG.0000000000001634

Scrucca, L., Fop, M., Murphy, T. B., & Raftery, A. E. (2016). mclust 5: Clustering, classification and density estimation using Gaussian finite mixture models. The R Journal, 8(1), 289–317.

Svedlund, J., Sjödin, I., & Dotevall, G. (1988). GSRS–a clinical rating scale for gastrointestinal symptoms in patients with irritable bowel syndrome and peptic ulcer disease. Dig. Dis. Sci., 33(2), 129–134.

Ze, X., Duncan, S. H., Louis, P., & Flint, H. J. (2012). Ruminococcus bromii is a keystone species for the degradation of resistant starch in the human colon. The ISME Journal, 6(8), 1535–1543. 10.1038/ismej.2012.4

